# First evidence that intrinsic fetal heart rate variability exists and is affected by hypoxic pregnancy

**DOI:** 10.1101/242107

**Authors:** Martin G. Frasch, Christophe L. Herry, Youguo Niu, Dino A. Giussani

## Abstract

Fetal heart rate variability (FHRV) emerges from influences of the autonomic nervous system, fetal body and breathing movements, and from baroreflex and circadian processes. We tested whether intrinsic HRV exists in the fetal period and whether it is affected by chronic fetal hypoxia. Chronically catheterized ewes carrying male singleton fetuses were exposed to normoxia (n=6) or hypoxia (10% inspired O2, n=9) for the last third of gestation (105-138 dG; term~145 dG) in isobaric chambers. At 138dG, isolated hearts were studied using a Langendorff preparation. We calculated basal iFHRV matrix indices across five signal-analytical domains from the systolic peaks within 15 min segments in each heart. Significance was assumed at p<0.05. Maternal hypoxia yielded chronic fetal PaO2 values of 11.5±0.6 (mean+SEM) relative to controls of 20.9±0.5 mmHg. Hearts of fetuses isolated from hypoxic pregnancy showed approximately 4-fold increases in the Grid transformation as well as the AND similarity index (sgridAND, informational domain) and a 4-fold reduction in the Scale dependent Lyapunov exponent slope (invariant domain). We also detected a 2-fold reduction in the Recurrence quantification analysis, percentage of laminarity and recurrences and maximum diagonal line (dlmax, geometric domain) and in the Multiscale time irreversibility asymmetry index (energetic domain). dlmax and sgridAND correlated with left ventricular end-diastolic pressure across both groups (R^2^=0.32 and R^2^=0.63, respectively). This is the first evidence that iHRV originates in fetal life and that chronic fetal hypoxia significantly alters it. Isolated fetal hearts from hypoxic pregnancy exhibit a lower complexity in iFHRV.

**Key points summary:**

- We introduce a technique to test whether intrinsic fetal heart rate variability (iFHRV) exists and we show the utility of the technique by testing the hypothesis that iFHRV is affected by chronic fetal hypoxia, one of the most common adverse outcomes of human pregnancy complicated by fetal growth restriction.
- Using an established late gestation ovine model of fetal development under chronic hypoxic conditions, we identify iFHRV in isolated fetal hearts and show that it is markedly affected by hypoxic pregnancy.
- Therefore, the isolated fetal heart carries a memory of adverse intrauterine conditions experienced during the last third of pregnancy.
- Indices of iFHRV may improve fetal health surveillance by helping to diagnose prediction of fetal hypoxia and acidaemia during antenatal or intra-partum fetal compromise.

## Introduction

Antenatal electronic monitoring of fetal heart rate (FHR) is widely used clinically and it is an important tool to assess the fetal condition (Westgate *et al.*, 2007). FHR monitoring is routinely used to assess fetal well-being in pregnancies affected by utero-placental dysfunction and profound, prolonged alterations in FHR variability (FHRV) are thought to represent acute fetal compromise (Yamaguchi *et al.*, 2018). Persistent reductions in short term variation (STV) below 3 ms in the antenatal period, within 24 hours of delivery, is moderately predictive of an increased risk of metabolic acidosis in the neonate, and early infant death (Infant Collaborative Group, 2017)(Kapaya *et al.*, 2016). As such, STV, as part of computerized cardiotocography (cCTG) examination, is not recommended as the sole means of antenatal surveillance of human fetuses with suspected utero-placental dysfunction (Grivell *et al.*, 2010). Therefore, there is an urgent need for additional indices of FHRV for diagnostic prediction of fetal hypoxia and acidaemia during antenatal or intra-partum fetal compromise.

It is established that normal FHRV represents a complex, nonlinear integration of the activities of the sympathetic and parasympathetic nervous systems. Fetal body and breathing movements, sleep states (Nijhuis *et al.*, 1982) as well as baroreflex and circadian processes also influence FHRV (Dalton *et al.*, 1977; Visser *et al.*, 1982; Frasch *et al.*, 2009; Jensen *et al.*, 2009). There is some evidence for intrinsic pacemaker rhythms of the sino-atrial node that affect HRV in critically ill adult patients (Papaioannou *et al.*, 2013). However, whether intrinsic HRV (iHRV) exists in the fetal period and whether it is affected by adverse intrauterine conditions has never been tested.

The isolated Langendorff *ex vivo* preparation of the fetal sheep heart is perfectly suited for assessing iHRV because it is devoid of any innervation or systemic hormonal influences. Therefore, the objectives of this work were to introduce to the field a new technique for physiological research and to show its utility by assessing whether iFHRV exists. Further, combining novel technology only recently available to induce chronic fetal hypoxia and fetal growth restriction in ovine pregnancy (Brain *et al.*, 2015a; Allison *et al.*, 2016), we tested whether iFHRV is affected in the compromised IUGR fetus in late gestation. The data show that iFHRV exists and that it is affected by chronic fetal hypoxia, thereby expanding technology to diagnose the chronically compromised fetus and improve fetal health surveillance.

## Methods

### Surgical Preparation

All experiments were performed in accordance with the UK Home Office guidance under the Animals (Scientific Procedures) Act 1986 and were approved by the Ethical Review Board of the University of Cambridge.

Briefly, chronically catheterized ewes carrying male singleton fetuses were exposed to normoxia (n=6) or hypoxia (10% inspired O2, n=9) for the last third of gestation (105-138 dG; term~145 dG) in bespoke isobaric chambers, (Brain *et al.*, 2015b; Allison *et al.*, 2016; Shaw *et al.*, 2018). At 138dG, isolated hearts were studied under a Langendorff preparation using established techniques (Fletcher *et al.*, 2005; Niu *et al.*, 2013, 2018) (Fig. 1).

**Figure 1.**
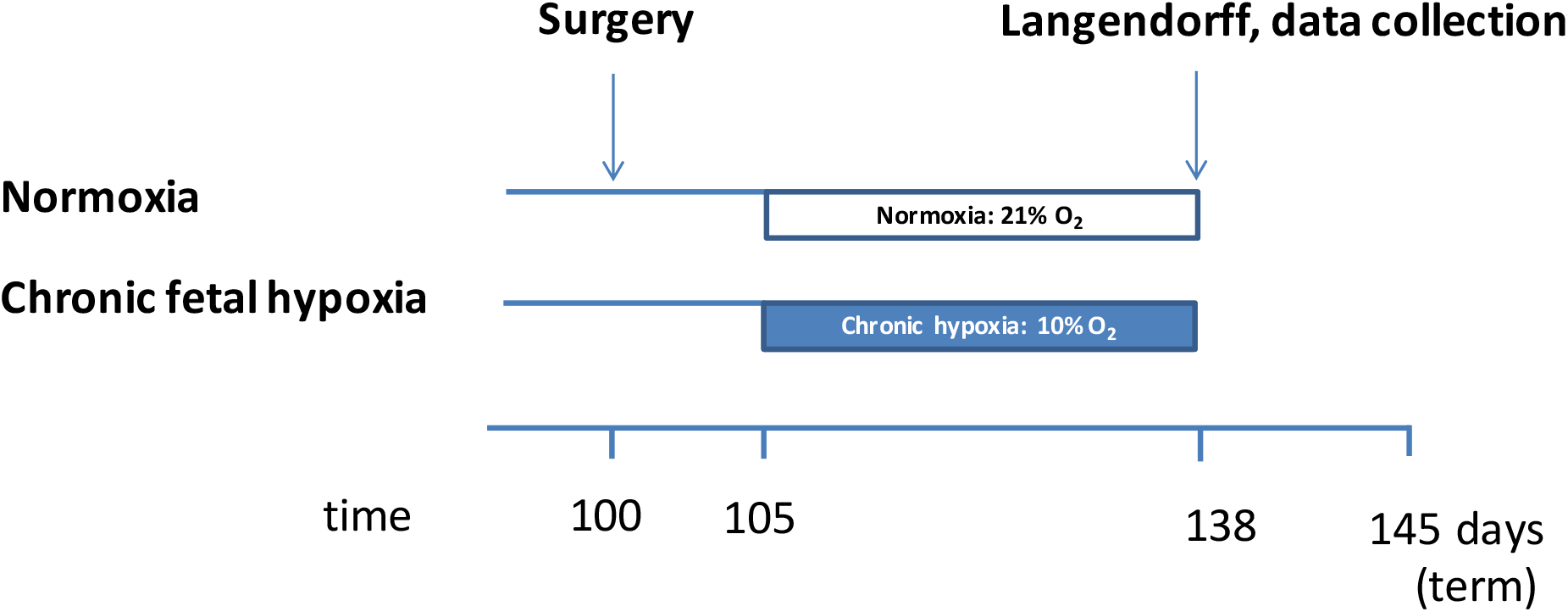
**LEFT**: Experimental protocol for *ex vivo* analyses. **MIDDLE**: Isobaric hypoxic chambers and nitrogen-generating system. **RIGHT**: Isolated heart perfusion model.

At 100±1 days gestational age (term ca. 145 days), pregnant Welsh mountain ewes carrying singleton pregnancies determined by ultrasound scan (Toshiba Medical Systems Europe, Zoetermeer, the Netherlands) underwent a laparotomy under general anaesthesia. In brief, food but not water was withdrawn for 24 h prior to surgery. Anaesthesia was induced by Alfaxan (1.5–2.5 mg kg1 i.v. alfaxalone; Jurox Ltd., Worcestershire, UK) and general anaesthesia (1.5–2.0% isoflurane in 60:40 O2:N2O) maintained by use of a positive pressure ventilator (Datex-Ohmeda Ltd., Hatfield, Hertfordshire, UK). Antibiotics (30 mg kg1 i.m. procaine benzylpenicillin; Depocillin; Intervet UK Ltd., Milton Keynes, UK) and an analgesic (1.4 mg kg1 s.c. carprofen; Rimadyl; Pfizer Ltd., Kent, UK) were administered immediately before the start of surgery. Following a midline abdominal incision and uterotomy, the fetal hind limbs were exposed, and the fetal sex was determined. If male, then the fetuses were chosen for this study. Female fetuses were used for another experiment. The fetus was returned into the intrauterine cavity, and the uterine and maternal abdominal incisions were closed in layers. A Teflon catheter (i.d. 1.0 mm, o.d. 1.6 mm, Altec, UK) was then placed in the maternal femoral artery and extended to the descending aorta, in addition to a venous catheter extended into the maternal inferior vena cava (i.d. 0.86 mm, o.d. 1.52 mm, Critchly Electrical Products, NSW, Australia). Catheters were filled with heparinized saline (80 I.U mL1 heparin in 0.9% NaCl), tunnelled subcutaneously, and exteriorized via a keyhole incision made in the maternal flank to be kept inside a plastic pouch sewn onto the maternal skin. Inhalation anaesthesia was withdrawn, and the ewe was ventilated until respiratory movements were observed. The ewe was extubated when spontaneous breathing returned and moved into a recovery pen adjacent to other sheep with free access to food and water. A total of 18 Welsh Mountain ewes carrying male singleton fetuses were surgically instrumented for this study.

### Postoperative care

Following surgery, ewes were housed in individual floor pens with a 12 h:12 h light:dark cycle and free access to hay and water. Antibiotics (30 mg kg1 i.m. procaine benzylpenicillin; Depocillin; Intervet UK Ltd., Milton Keynes, UK) were administered daily to the ewe for 5 days following surgery. From 103 days of gestation, ewes were fed daily a bespoke maintenance diet made up of concentrate and hay pellets to facilitate the monitoring of food intake (Cambridge ewe diet: 40 g nuts kg1 and 3 g hay kg1; Manor Farm Feeds Ltd.; Oakham, Leicestershire, UK). Generally, normal feeding patterns were restored within 24–48 h of recovery. On day 103 of gestation, ewes were randomly assigned to either of two experimental groups: normoxia (N: n = 6) or chronic hypoxia (H: n = 9).

Ewes allocated to chronic hypoxic pregnancy were housed in one of four bespoke isobaric hypoxic chambers (Telstar Ace, Dewsbury, West Yorkshire, UK; Fig. 1), as previously described (Brain *et al.*, 2015a; Allison *et al.*, 2016). In brief, chambers were supplied with variable amounts of nitrogen and air provided via nitrogen generators and air compressors, respectively, from a custom-designed nitrogen-generating system (Domnick Hunter Gas Generation, Gateshead, Tyne & Wear, UK). Ambient PO2, PCO2, humidity, and temperature within each chamber were monitored via sensors, displayed, and values recorded continuously via the Trends Building Management System of the University of Cambridge through a secure Redcare intranet.

In this way, the percentage of oxygen in the isolators could be controlled with precision continuously over long periods of time. For experimental procedures, each chamber had a double transfer port to internalize material and a manually operated sliding panel to encourage the ewe into a position where daily sampling of blood could be achieved through glove compartments (Fig. 1). Pregnancies assigned to the chronic hypoxia group were placed inside the chambers at 103 days of gestation under normoxic conditions (11 L sec-1 air, equating to 39.6 m3 h-1). At 105 days, pregnancies were exposed to approximately 10% O2 by altering the incoming inspirate mixture to 5 L sec1 air: 6 L sec1 N2.

### Langendorff preparation

Fetal hearts were isolated, mounted onto a Langendorff apparatus and perfused at a constant pressure of 30 mmHg, as detailed by (Fletcher *et al.*, 2005). The ductus arteriosus was ligated. Pulmonary arteriotomy was performed. A recirculating solution of Krebs-Henseleit bicarbonate buffer containing (mM.L^−1^) 120 NaCl, 4.7 KCl, 1.2 MgSO_2_.7H_2_O, 1.2 KH_2_PO_4_, 25 NaHCO_3_, 10 glucose, and 1.3 CaCl_2_.2H_2_O was filtered through a 5 μm cellulose nitrate filter (Millipore, Bedford, MA, USA) and gassed with O_2_:CO_2_ (95:5) at 37°C. A small flexible non-elastic balloon was inserted into the left ventricle through the left atrium. The balloon was filled with deionised water and attached to a rigid deionised water-filled catheter connected to a calibrated pressure transducer (Argon Medical Devices, Texas, USA). The balloon volume was adjusted to obtain a left ventricular end diastolic pressure (LVEDP) recording of approximately 5-10 mmHg. After an initial 15 min stabilisation period, basal heart rate (HR), left ventricular systolic pressure (LVSP) and LVEDP were recorded. Basal left ventricular developed pressure (LVDP) was calculated as LVSP-LVEDP. The maximum and minimum first derivatives of the left ventricular pressure (dP/d*t*_max_ and dP/d*t*_min_) were calculated using an M-PAQ data acquisition system (Maastricht Programmable AcQuisition System, Netherlands). For HRV analysis purpose, all original recording traces of left ventricular pressure were exported to LabChart^®^ 7 software (ADInstruments, UK).

### FHRV analysis

To derive FHRV, recordings of fetal left ventricular pressure sampled at 1kHz were analyzed with the CIMVA (continuous individualized multiorgan variability analysis) software, as before (Durosier *et al.*, 2014). Inter-beat intervals were extracted from the pressure recordings using the systolic peaks. A range of 55 basal iHRV indices was then calculated across five signal-analytical domains from the inter-beat interval time series, within 15 min segments in each heart, determined as an average of three non-overlapping 5 min intervals.

### Statistical analysis

Data are presented as Mean+SEM. The Student’s *t* test for unpaired data was used to compare variables from hypoxic versus normoxic pregnancy. Relationship between variables were assessed by the Spearman rank correlation. Statistical significance was set at P<0.05 (SigmaStat).

## Results

Maternal P_a_O_2_ was reduced in hypoxic pregnancy (42.0±1.2 vs. 105.7±3.7 mmHg, P<0.05). Fetuses exposed to chronic hypoxia had a significant reduction in the partial pressure of arterial oxygen in the descending aorta (11.5±0.6 vs. 20.9±0.5mmHg, P<0.05).

Hearts isolated from chronically hypoxic fetuses showed distinct changes in iHRV measures across four signal-analytical domains (Fig. 2). There were approximately 4-fold increases in the Grid transformation feature as well as the AND similarity index (sgridAND, informational domain) and a 4-fold reduction in the Scale dependent Lyapunov exponent slope (SDLEalpha, invariant domain). We also detected a 2-fold reduction in the Recurrence quantification analysis, the percentage of laminarity and recurrences and maximum diagonal line (pL, pR, dlmax, all from geometric domain) and in the Multiscale time irreversibility asymmetry index (AsymI, energetic domain). There was also a moderate fall in the Detrended fluctuation analysis (area under the curve, DFA AUC, invariant domain).

**Figure 2.**
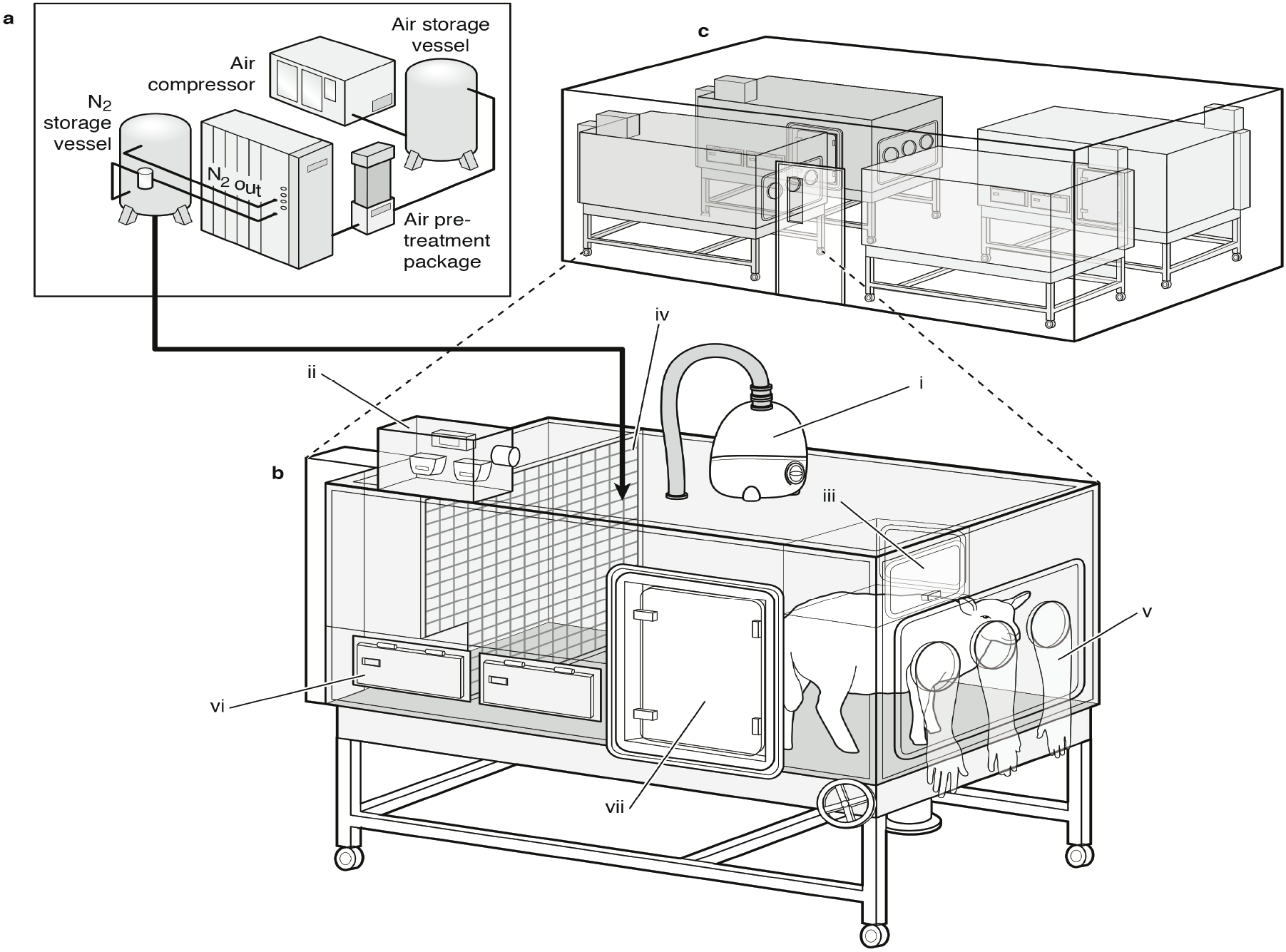
Effects of chronic hypoxia during pregnancy on fetal intrinsic heart rate variability (iFHRV). Values are mean ± SEM. Groups are normoxic (N, n=6) and hypoxic (H, n=9). Significant differences are: * vs. N, P<0.05 (Student’s t test for unpaired data).

Combined, these data suggest that isolated fetal hearts from control pregnancy exhibited significant intrinsic FHRV. Further, isolated fetal hearts from hypoxic pregnancy showed a significantly lower complexity in iFHRV. Measures of dlmax and sgridAND also correlated with left ventricular end-diastolic pressure (LVEDP) across both groups (Fig. 3).

**Figure 3.**
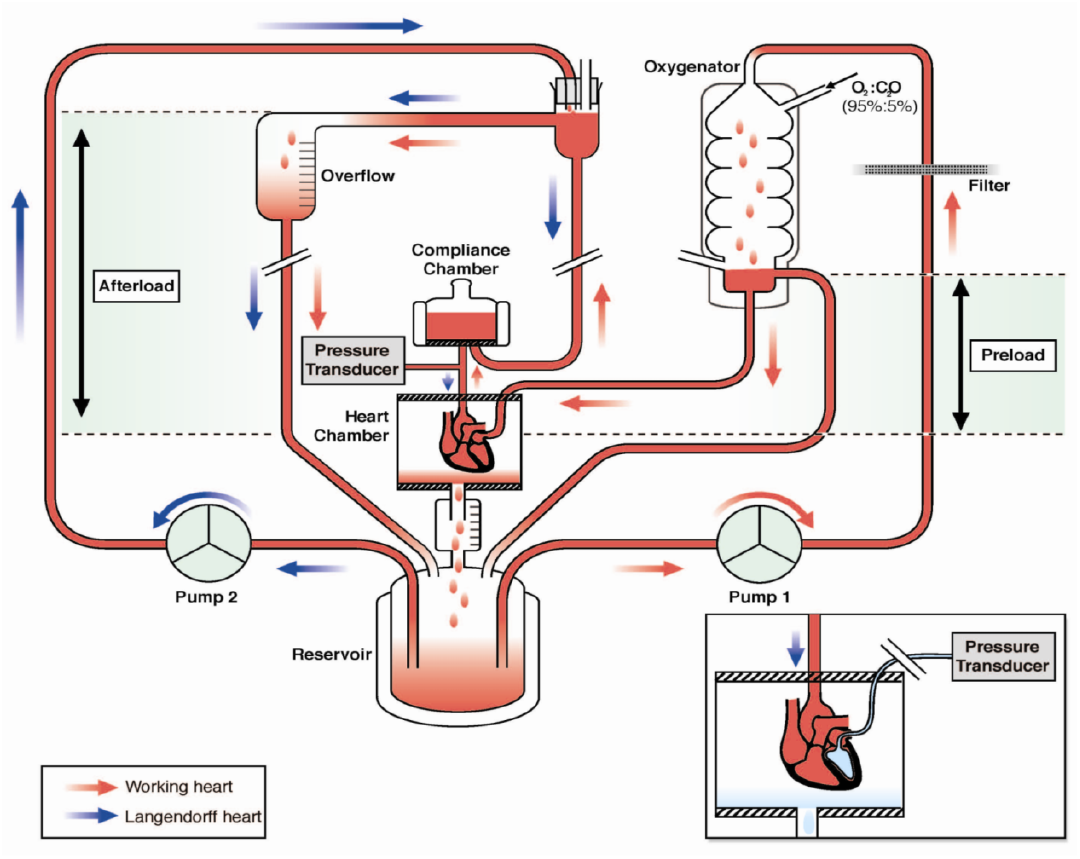
Correlation of left ventricular end-diastolic pressure (LVEDP) and the intrinsic heart rate variability (iHRV) measures dlmax and sgridAND. LVEDP in normoxic (empty circles) and hypoxic (black circles) groups. Spearman statistics: R^2^=0.32, p=0.03, and R^2^=0.63, p<0.001, respectively.

## Discussion

The data show that the fetal heart in late gestation has intrinsic influences, which may affect fetal cardiac function, and which can be picked up by FHR monitoring. Further, iFHRV is significantly affected by pregnancy complicated by chronic hypoxia. Combined, these discoveries provide a conceptual advance to this field of study, which may help in the interpretation of fetal heart rate patterns. Hence, the data provide potential translational biomarkers for the improved diagnosis of fetal health in human complicated pregnancy.

Analysis of FHRV has served as a scientific and diagnostic tool to quantify the fluctuations of cardiac activity under various conditions since the early 1980’s (Akselrod *et al.*, 1981). However, surprisingly, little is known about its biological origins. From studies of healthy adult subjects during exercise and investigations of heart-transplant recipients, the field is aware that intrinsic components of cardiac rhythm can contribute substantially to HRV (Akselrod *et al*., 1981). However, a prenatal origin of intrinsic influences in HRV has been difficult to prove. Further, if iFHRV occurs, whether it is affected by chronic fetal hypoxia, one of the most common outcomes of human pregnancy complicated by fetal growth restriction, is completely unknown.

Here, we provide the first evidence that iHRV originates in fetal life and that chronic fetal hypoxia significantly alters it. The significant relationship between nonlinear measures of FHRV and changes in left ventricular end diastolic pressure (LVEDP), which is elevated in fetuses from hypoxic pregnancy (Niu *et al.*, 2018) suggests that such FHRV measures may reflect fetal myocardial dysfunction, particularly during cardiac diastole. Therefore, alterations in iFHRV may prove clinically useful as a biomarker of impaired cardiac reserve and fetal myocardial decompensation during antepartum or intrapartum monitoring of the fetal heart in risky pregnancy or complicated labour.

The findings raise several questions. What are the mechanisms contributing to iFHRV in the late gestation fetus in normal healthy pregnancy? What is the transfer mechanism by which *in utero* chronic hypoxia imprints upon iFHRV? May it be via impacting on myocardiogenesis, which then affects patterns of cardiac contractility and relaxability, such as alterations in LVEDP? Does the putative transfer mechanism of *in utero* hypoxia upon iFHRV depend upon vagal and sympathetic fluctuations *in vivo* or is it entirely autochthonic, emerging from the adaptive processes within the excitatory cells themselves in response to chronic hypoxia?

Previous findings derived from sheep studies in which fetuses were subjected to a labour-like insult with worsening acidaemia and work in adult animal models of acidaemia indicate that around a pH of 7.2, the physiological myocardial activity is curbed via a Bezold-Jarisch-like reflex. This is a vagally-mediated myocardial depressive reflex that reduces cardiac output under conditions of moderate acidaemia, thereby preserving depleting myocardial energy reserves (Harry *et al.*, 1971; Nuwayhid *et al.*, 1975; Nuyt *et al.*, 2001; Frasch *et al.*, 2008; Gold *et al.*, 2017). When labour is associated with worsening fetal acidaemia, fetal compensatory cardiovascular reflexes are sensitized (Thakor & Giussani, 2009) and at risk of becoming overwhelmed, leading to eventual cardiac decompensation and an increased risk of fetal brain injury (Yumoto *et al.*, 2005). Fetal acidaemia impacts upon fetal myocardial contractility, which further promotes decreased cardiac output and the inability to maintain fetal arterial blood pressure (Frasch *et al.*, 2008, 2011). It is not yet understood how fetal insults involving hypoxia with or without worsening acidaemia disrupt sinus node pacemaker activity, thereby affecting iFHRV.

It is interesting to consider what the identified iHRV features dlmax and sgridAND, which correlate to LVEDP, may represent physiologically. The first measure is derived from a recurrence plot where the diagonal lines represent the trajectory visiting the same region of the phase space at different times. The lengths of diagonal lines in a recurrence plot are related to the predictability of the system dynamics. Perfectly predictable systems would have infinitely long diagonal lines in the recurrence plot (high dlmax). Conversely, stochastic and chaotic systems would have very short diagonal lines (low dlmax)(Webber & Zbilut, 1994; Zbilut *et al.*, 2002; Webber & Marwan, 2015). In the present study, chronic hypoxic pregnancy reduced the dlmax component of iFHRV, which correlated to an elevated LVEDP compared to hearts isolated from normoxic fetuses.

The grid transformation AND similarity index (sgridAND) measures the dynamic system phase space reconstruction trajectory, with a specific embedding dimension and time delay. It is binarized over a grid (i.e., pixel visited by the trajectory=1, all others=0) to produce an image. Two grid images corresponding to different time delays or different windows in time are then compared (Roopaei *et al.*, 2010). The sgridAND measure is the normalized sum of the binary AND operation on the two compared images and represents a similarity index between the phase space trajectory from two consecutive windows. Low values indicate that the iHRV dynamics have changed while high values mean the dynamics are similar. The latter is the case for the iFHRV calculated in hearts isolated from hypoxic fetuses (Fig. 4). Again, this correlates with greater resting LVEDP.

**Figure 4.**
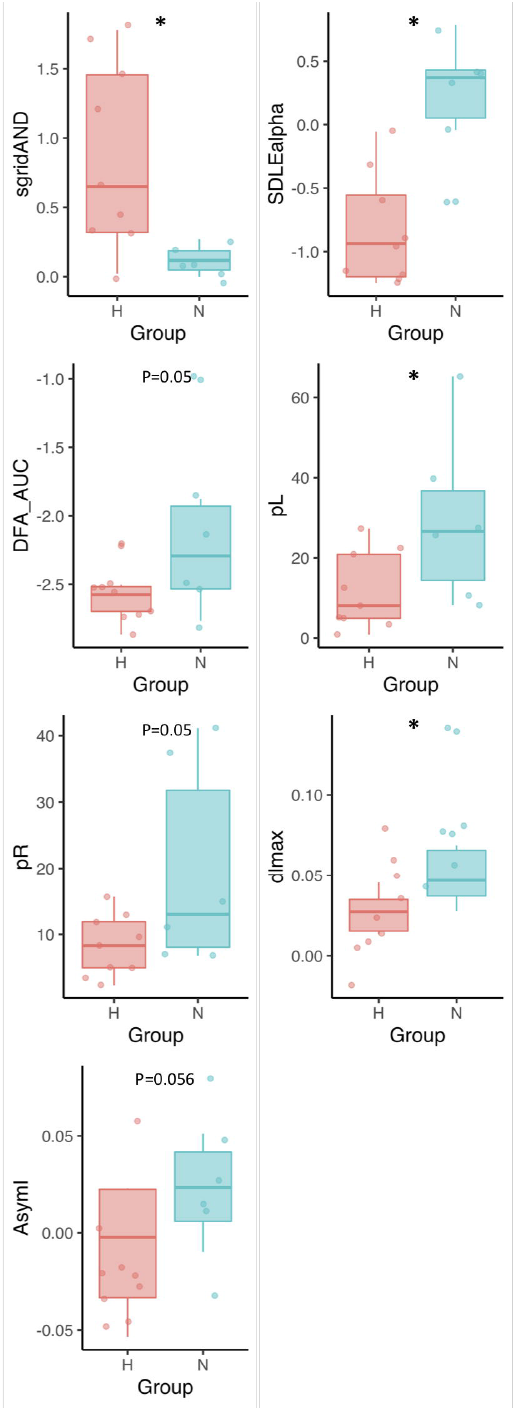
A representative beat-to-beat time series, followed by the corresponding grid transformation AND similarity index (sgridAND) and the recurrence plots are shown for a normoxic (right) and hypoxic (left) fetus to demonstrate iFHRV pattern differences revealed with such representation of the phase space organization of the beat-to-beat fluctuations.

Combined, our findings indicate that *in utero* hypoxia reduces the short-term predictability of iFHRV and increases its long-range similarity. Both effects do not contradict each other, because the effects are captured in different signal-analytical domains, one being a geometric feature of iFHRV and another referring to longer-term temporal processes in the informational domain. Importantly, both changes occur with a consistent increase in LVEDP, demonstrating that complex iFHRV properties can be linked to a cardiac phenotype.

### Study limitations

This investigation was conducted in an ovine *ex vivo* fetal heart preparation. Albeit derived in one of the most appropriate animal species that shares with humans similar temporal profiles of cardiac development (Morrison *et al.*, 2018), the present findings must be validated in human cohorts. This could be performed in the context of human heart transplants, which will likely require a multi-site effort, because at this age it is rare, with ~10 transplants performed in the US per year. (John & Bailey, 2018)

### Conclusions

We introduce a technique to the field of study that determines changes in iFHRV and validate the technique by showing that iFHRV measures can be significantly affected by chronic fetal hypoxia in physiological and clinically meaningful ways. These discoveries could lead to potential biomarkers that can be captured non-invasively from fetal ECG which may lead to improved diagnosis of fetal health antenatally and intra-partum. This may contribute to an improved and more timely diagnosis of fetal cardiovascular decompensation in human high risk pregnancy, before it is too late to prevent fetal brain injury (Gold *et al.*, 2017; Frasch, 2018; Yamaguchi *et al.*, 2018).

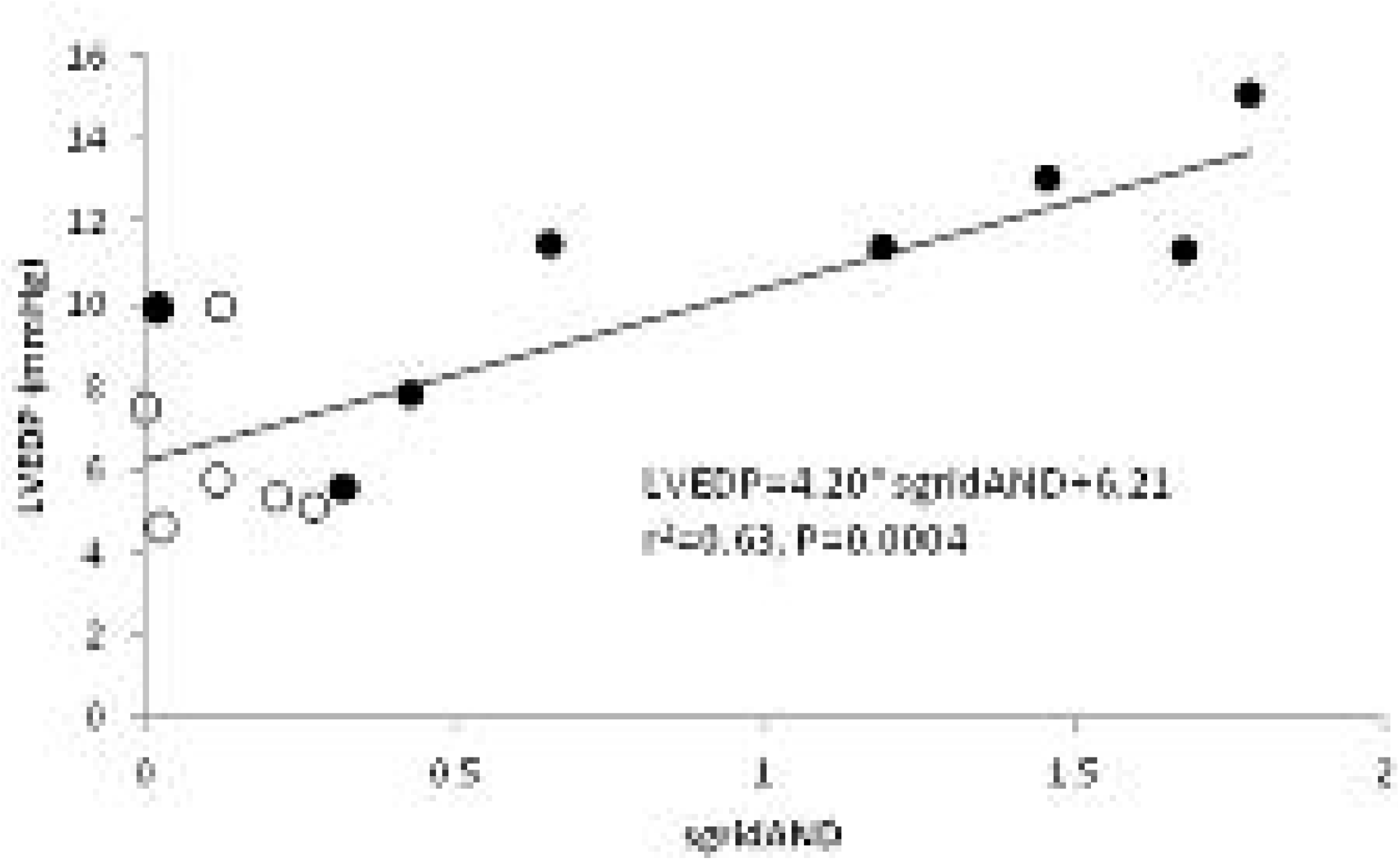

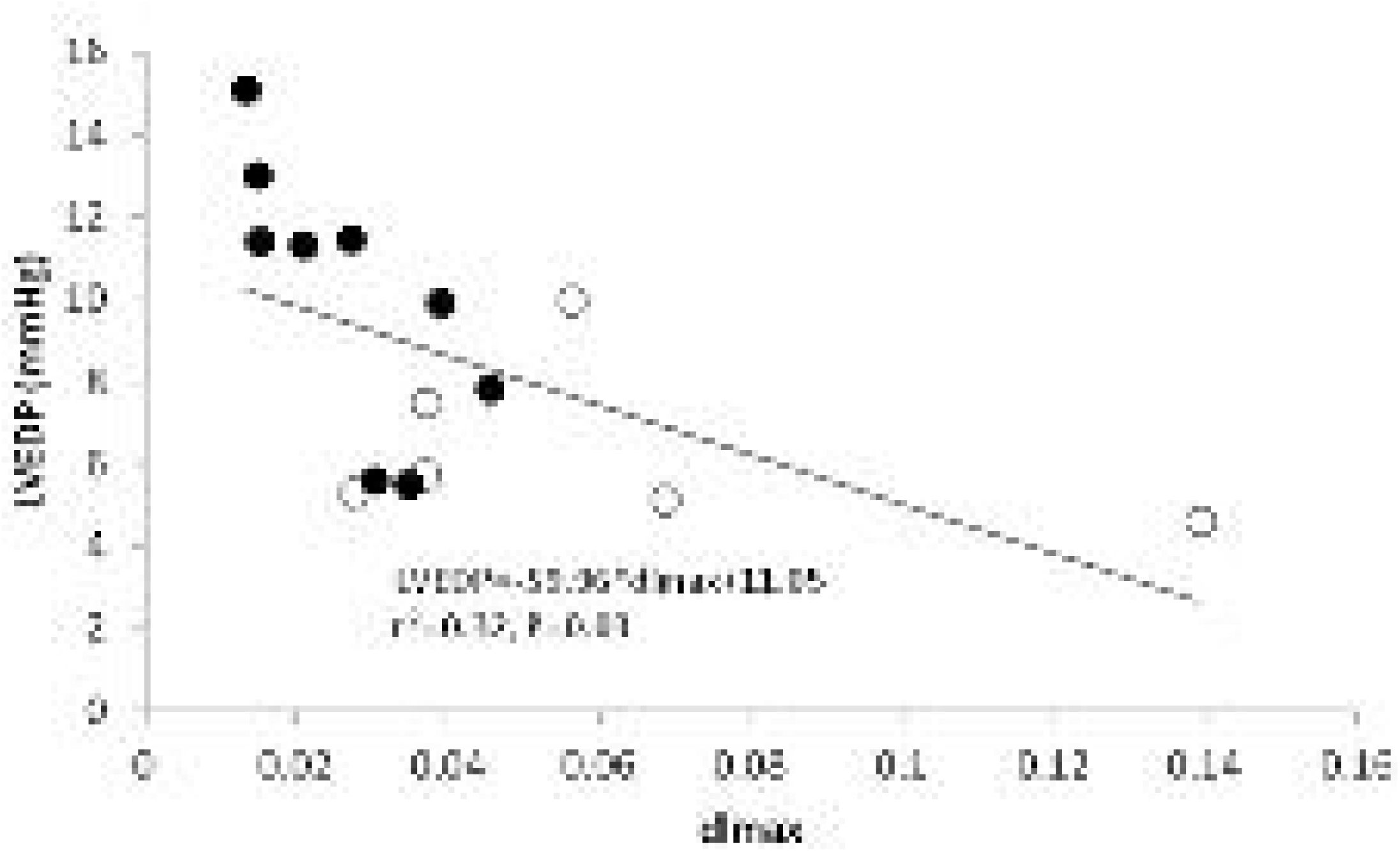

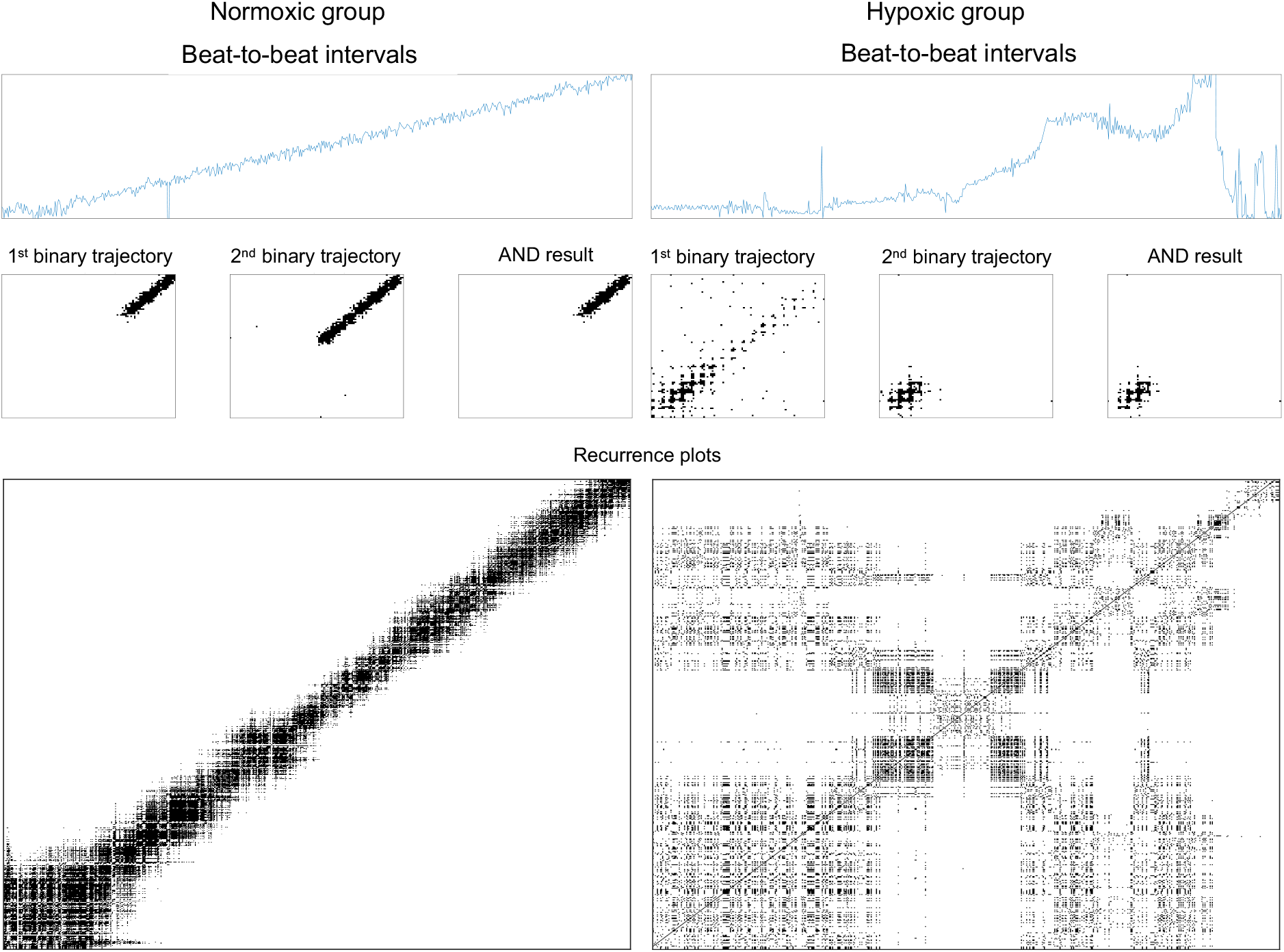

## Non-standard Abbreviations and Acronyms

CIMVA: continuous individualized multiorgan variability analysis;
FHR: fetal heart rate;
FHR variability: FHRV;
iFHRV: intrinsic FHRV;
LVEDP: left ventricular end-diastolic pressure;
sgridAND: grid transformation AND similarity index
IUGR: intrauterine growth restriction

## Acknowledgements

N/a.

## Sources of Funding

Supported by CIHR and The British Heart Foundation.

## Disclosures

The authors have nothing to disclose.

## References

Akselrod S, Gordon D, Ubel FA, Shannon DC, Berger AC & Cohen RJ (1981). Power spectrum analysis of heart rate fluctuation: a quantitative probe of beat-to-beat cardiovascular control. Science 213, 220–222.

Allison BJ, Brain KL, Niu Y, Kane AD, Herrera EA, Thakor AS, Botting KJ, Cross CM, Itani N, Skeffington KL, Beck C & Giussani DA (2016). Fetal in vivo continuous cardiovascular function during chronic hypoxia. J Physiol 594, 1247–1264.

Brain KL, Allison BJ, Niu Y, Cross CM, Itani N, Kane AD, Herrera EA & Giussani DA (2015a). Induction of controlled hypoxic pregnancy in large mammalian species. Physiol Rep; DOI:DOI 10.14814/phy2.12614.

Brain KL, Allison BJ, Niu Y, Cross CM, Itani N, Kane AD, Herrera EA & Giussani DA (2015b). Induction of controlled hypoxic pregnancy in large mammalian species. Physiol Rep; DOI:10.14814/phy2.12614.

Dalton KJ, Dawes GS & Patrick JE (1977). Diurnal, respiratory, and other rhythms of fetal heart rate in lambs. Am J Obstet Gynecol 127, 414–424.

Durosier LD, Green G, Batkin I, Seely AJ, Ross MG, Richardson BS & Frasch MG (2014). Sampling rate of heart rate variability impacts the ability to detect acidemia in ovine fetuses near-term. Front Pediatr 2, 38.

Fletcher AJW, Forhead AJ, Fowden AL, Ford WR, Nathanielsz PW & Giussani DA (2005). Effects of gestational age and cortisol treatment on ovine fetal heart function in a novel biventricular Langendorff preparation. J Physiol 562, 493–505.

Frasch MG (2018). Saving the brain one heartbeat at a time. J Physiol; DOI: 10.1113/JP275776.

Frasch MG, Keen AE, Gagnon R, Ross MG & Richardson BS (2011). Monitoring fetal electrocortical activity during labour for predicting worsening acidemia: a prospective study in the ovine fetus near term. PLoS One 6, e22100.

Frasch MG, Mansano R, Ross MG, Gagnon R & Richardson BS (2008). Do repetitive umbilical cord occlusions (UCO) with worsening acidemia induce the Bezold-Jarisch reflex (BJR) in the ovine fetus near term? Reprod Sci 15, 129A.

Frasch MG, Muller T, Hoyer D, Weiss C, Schubert H & Schwab M (2009). Nonlinear properties of vagal and sympathetic modulations of heart rate variability in ovine fetus near term. Am J Physiol Regul Integr Comp Physiol 296, R702–R707.

Gold N, Frasch MG, Herry CL, Richardson BS & Wang X (2017). A Doubly Stochastic Change Point Detection Algorithm for Noisy Biological Signals. Front Physiol 8, 1112.

Grivell RM, Alfirevic Z, Gyte GM & Devane D (2010). Antenatal cardiotocography for fetal assessment. Cochrane Database Syst Rev 2010/01/22, CD007863.

Harry JD, Kappagoda CT, Linden RJ & Snow HM (1971). Depression of the reflex tachycardia from the left atrial receptors by acidaemia. J Physiol 218, 465–475.

Infant Collaborative Group (2017). Computerised interpretation of fetal heart rate during labour(INFANT): a randomised controlled trial. Lancet 389, 1719–1729.

Jensen EC, Bennet L, Guild SJ, Booth LC, Stewart J & Gunn AJ (2009). The role of the neural sympathetic and parasympathetic systems in diurnal and sleep state-related cardiovascular rhythms in the late-gestation ovine fetus. Am J Physiol Regul Integr Comp Physiol 297, R998–R1008.

John M & Bailey LL (2018). Neonatal heart transplantation. Ann Cardiothorac Surg 7, 118–125.

Kapaya H, Jacques R, Rahaim N & Anumba D (2016). “Does short-term variation in fetal heart rate predict fetal acidaemia?” A systematic review and meta-analysis. J Matern Fetal Neonatal Med 29, 4070–4077.

Morrison JL et al. (2018). Improving pregnancy outcomes in humans through studies in sheep. Am J Physiol Regul Integr Comp Physiol; DOI: 10.1152/ajpregu.00391.2017.

Nijhuis JG, Prechtl HF, Martin CB Jr & Bots RS (1982). Are there behavioural states in the human fetus? Early Hum Dev 6, 177–195.

Niu Y, Herrera EA, Evans RD & Giussani DA (2013). Antioxidant treatment improves neonatal survival and prevents impaired cardiac function at adulthood following neonatal glucocorticoid therapy. J Physiol 591, 5083–5093.

Niu Y, Kane AD, Lusby CM, Allison BJ, Chua YY, Kaandorp JJ, Nevin-Dolan R, Ashmore TJ, Blackmore HL, Derks JB, Ozanne SE & Giussani DA (2018). Maternal Allopurinol Prevents Cardiac Dysfunction in Adult Male Offspring Programmed by Chronic Hypoxia During Pregnancy. Hypertension 72, 971–978.

Nuwayhid B, Brinkman CR 3rd, Su C, Bevan JA & Assali NS (1975). Development of autonomic control of fetal circulation. Am J Physiol 228, 337–344.

Nuyt AM, Segar JL, Holley AT & Robillard JE (2001). Autonomic adjustments to severe hypotension in fetal and neonatal sheep. Pediatr Res 49, 56–62.

Papaioannou VE, Verkerk AO, Amin AS & de Bakker JM (2013). Intracardiac origin of heart rate variability, pacemaker funny current and their possible association with critical illness. Curr Cardiol Rev 9, 82–96.

Roopaei M, Boostani R, Sarvestani RR, Taghavi MA & Azimifar Z (2010). Chaotic based reconstructed phase space features for detecting ventricular fibrillation. Biomed Signal Process Control 5, 318–327.

Shaw CJ, Allison BJ, Itani N, Botting KJ, Niu Y, Lees CC & Giussani DA (2018). Altered autonomic control of heart rate variability in the chronically hypoxic fetus. J Physiol 596, 6105–6119.

Thakor AS & Giussani DA (2009). Effects of acute acidemia on the fetal cardiovascular defense to acute hypoxemia. American journal of physiology Regulatory, integrative and comparative physiology 296, R90–R99.

Visser GH, Goodman JD, Levine DH & Dawes GS (1982). Diurnal and other cyclic variations in human fetal heart rate near term. Am J Obstet Gynecol 142, 535–544.

Webber CL Jr & Marwan N (2015). Recurrence quantification analysis. Springer.

Webber CL Jr & Zbilut JP (1994). Dynamical assessment of physiological systems and states using recurrence plot strategies. J Appl Physiol 76, 965–973.

Westgate JA, Wibbens B, Bennet L, Wassink G, Parer JT & Gunn AJ (2007). The intrapartum deceleration in center stage: a physiologic approach to the interpretation of fetal heart rate changes in labor. Am J Obstet Gynecol 197, 236.e1–e11.

Yamaguchi K, Lear CA, Beacom MJ, Ikeda T, Gunn AJ & Bennet L (2018). Evolving changes in fetal heart rate variability and brain injury after hypoxia-ischaemia in preterm fetal sheep. J Physiol; DOI: 10.1113/JP275434.

Yumoto Y, Satoh S, Fujita Y, Koga T, Kinukawa N & Nakano H (2005). Noninvasive measurement of isovolumetric contraction time during hypoxemia and acidemia: Fetal lamb validation as an index of cardiac contractility. Early Hum Dev 81, 635–642.

Zbilut JP, Thomasson N & Webber CL (2002). Recurrence quantification analysis as a tool for nonlinear exploration of nonstationary cardiac signals. Med Eng Phys 24, 53–60.

